# Cholesterol catalyzes unfolding in membrane inserted motifs of the pore forming protein cytolysin A

**DOI:** 10.1101/2023.05.07.539733

**Authors:** Avijeet Kulshrestha, Sudeep N Punnathanam, Rahul Roy, K Ganapathy Ayappa

**Affiliations:** Department of Chemical Engineering, Indian Institute of Science, Bangalore, India - 560012; Center for BioSystems Science and Engineering, Indian Institute of Science, Bangalore, Karnataka, India - 560012

## Abstract

Plasma membrane induced protein folding and conformational transitions play a central role in cellular homeostasis. Several transmembrane proteins are folded in the complex lipid milieu to acquire a specific structure and function. Bacterial pore forming toxins (PFTs) are proteins expressed by a large class of pathogenic bacteria that exploit the plasma membrane environment to efficiently undergo secondary structure changes, oligomerize and form transmembrane pores. Unregulated pore formation causes ion imbalance leading to cell death and infection. Determining the free energy landscape of these membrane driven transitions remains a challenging problem. Although cholesterol recognition is required for lytic activity of several proteins in the PFT family of toxins, the regulatory role of cholesterol for the *α*-PFT, cytolysin A expressed by E. coli is less understood. In a recent free energy computation, we have shown that the *β*-tongue, a critical membrane inserted motif of the ClyA toxin, has an on-pathway partially unfolded intermediate that refolds into the helix-turn-helix motif of the pore state.^1^ To understand the molecular role played by cholesterol, we have carried out string method based computations in membranes devoid of cholesterol which reveals an increase of *∼* 30 times in the free energy barrier for the loss of *β*-sheet secondary structure when compared with membranes containing cholesterol. Specifically the tyrosine-cholesterol interaction was found to be critical to stabilizing the unfolded intermediate. In the absence of cholesterol the membrane was found to undergo large curvature deformations in both leaflets of the membrane accompanied by bilayer thinning. Our study with the *α*-toxin, ClyA illustrates that cholesterol is critical to catalyzing and stabilizing the unfolded state of the *β*-tongue in the membrane, opening up fresh insights into cholesterol assisted unfolding of membrane proteins.

**Significance:** Cholesterol, an integral part of mammalian cell membranes, is necessary for activity of pathogenic toxins. Our understanding of the thermodynamic and molecular underpinnings of cholesterol-protein interactions during different stages of toxin activity is unclear. Using path based all atom molecular dynamics simulations, we illustrate lowered free energy barriers and enhanced stability of the membrane unfolded intermediate of an *α*-pore forming toxin (PFT) ‘ClyA’ providing insights into the increased pore formation kinetics with cholesterol. Thus, membrane cholesterol generally believed to play a passive receptor function for PFT activity is involved in a more complex regulatory role in assisting secondary structure transitions critical to PFT lytic activity. Our findings could aid in drug development strategies for mitigating PFT mediated bacterial infections.

## Introduction

The environment of the plasma membrane plays a vital regulatory role in stabilizing membrane proteins important for cellular homeostasis. ^2^ Transmembrane proteins are predominantly *α*-helical in structure and include the G-protein family of receptors (GPCRs), ion channels, and aquaporins.^3^ The translocon mediated mechanism for protein insertion and folding in the plasma membrane has been widely investigated, and provides a cotranslational pathway for protein synthesis and refolding in the phospholipid membrane.^4^ In contrast, pathogenic bacteria have evolved to express pore forming proteins (PFTs) that form large transmembrane pore complexes on the plasma membrane using efficient membrane binding, as well as unfolding and refolding events of membrane inserted motifs.^1, 5, 6^ The process of oligomerization to form the oligomeric pore complex is often accompanied by large conformational changes that occur at the lipid interface. Since conformational changes and folding occur spontaneously in the membrane, studying the transition of pore forming proteins from a water soluble monomeric state to the membrane inserted protomer is expected to provide several insights into membrane driven folding of proteins.

PFTs are a pathogenic class of the family of pore-forming proteins, which are expressed by bacteria to mount infections mediated by membrane assisted assembly and pore formation.^6^ Pore formation allows the passage of crucial cellular components of different sizes ranging from ions to ATP molecules leading to cell death. Pores also provide passage to other proteins known as lethal factors to promote infections. PFTs are classified based on the secondary structure of the transmembrane region, which is *α*-helical for the *α*-PFTs and *β*-barrels for the *β*-PFTs.^5^ In this study, we focus on cytolysin A (ClyA), expressed by *Escherichia coli*, *Salmonella typhi*, and *Shigella flexneri*.^6^ ClyA is a 34 kDa *α*-PFT which shows one of the largest conformational changes in PFTs, oligomerizing into a dodecameric pore state on the membrane. ClyA has more recently been the subject of investigation because of the availability of high resolution crystal structures^7, 8^ of the monomer (Figure 1A) and the protomer states (Figure 1B). The major conformational changes that occur in ClyA during the transition from the monomer to the protomer states are the flipping out of the N-terminus from the *α*-helical bundle and the conversion of *β*-tongue to the helix-turn-helix motif as illustrated in Figure 1. Using molecular dynamics (MD) simulations with structure based models (SBMs) the monomer to protomer transition was shown to be triggered by the interaction of the predominantly hydrophobic *β*-tongue motif with the membrane,^9^ confirming the original hypothesis postulated during the crystal structure elucidation of ClyA.^8^

**Figure 1:**
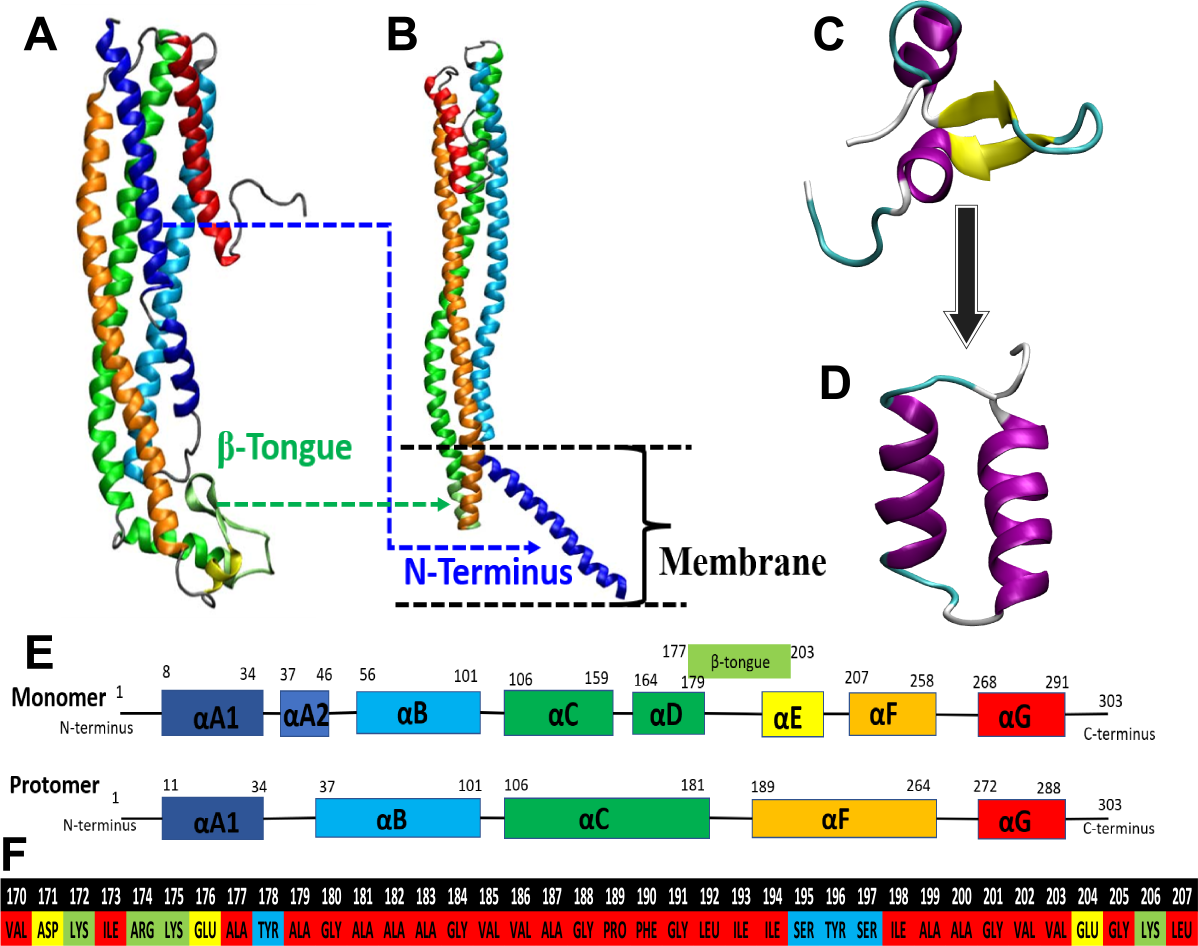
Cytolysin A structure detail. (A) Water-soluble monomer structure, (B) membrane inserted protomer structure, (C) *β*-tongue structure (170-207), (D) helix-turn-helix motif (170-207) of protomer state, (E) secondary structure detail of monomer and the protomer states, (F) sequence of the amino acid residues of the truncated *β*-tongue region used in free energy computations - Red: Non-polar, Blue: Polar, Green: positively charged, Yellow: Negatively charged. Adapted from Kulshrestha et al. ^1^, Copyright 2023, American Chemical Society.

Using string method based free energy computations ^10^ and MD simulations in a POPC:Chol (70:30) membrane, we have recently shown^1^ that the transition of the membrane inserted *β*-tongue to the helix-turn-helix motif of the protomer state takes places through a partially unfolded intermediate. This transition to the unfolded state is spontaneous and the refolding to the helix-turn-helix motif was shown to have a barrier of *∼* 26.56 kJ/mol. We further showed that single point mutations that lowered the flexibility of the *β*-tongue motif resulted in impaired pore formation kinetics indicating that the inherent flexibility was critical to membrane induced re-folding and effective pore formation. Cholesterol is a component of the mammalian cell membrane and several PFTs, such as listeriolysin O (LLO) and pneumolysin (PLY), which belong to the class of cholesterol-dependent cytolysins (CDCs), code specificity into targeting cells by using cholesterol as a receptor.^11–13^ Although ClyA is not part of the CDC family of toxins, it has been shown to have higher pore forming and lytic activity in the presence of cholesterol. ^14, 15^ In a combined single molecule and simulation study in our laboratory we showed that cholesterol plays a significant role in enhancing the pore forming activity of ClyA^16^ and stabilized the pore complex by binding into pockets formed by adjacent *β*-tongues of the pore complex. Whether cholesterol plays a role in membrane binding and/or assists the refolding of the *β*-tongue is an important question with broad implications for other membrane protein folding transitions. In order to obtain a deeper understanding of the cholesterol role, we study the nature of the folding intermediate and its presence in membranes devoid of cholesterol and make a detailed comparison with free energy computations reported for membranes containing cholesterol.^1^

Computing the free energy of the monomer conformational changes, initiated by the membrane interaction, for a large 34 kDa protein of ClyA is extremely challenging and would require a judicious choice of collective variables to enable reliable sampling of the free energy landscape. We focus only on the key membrane binding *β*-tongue motif of the protein and compute the free energy associated with this transition in the absence of cholesterol. A detailed analysis of the trajectories along the path are carried out to determine the pivotal role played by cholesterol. We applied the finite-temperature string (FTS) method^1, 10^ in combination with path collective variables (PCVs)^17^ to obtain the transition path from the monomer state to the protomer state. Our study reveals a large free energy barrier associated with the unfolding transition of the *β*-tongue in a POPC membrane devoid of cholesterol. This barrier is about 30 times greater than the barrier observed with cholesterol where partial unfolding occurred spontaneously. Tyrosine known to be associated with anchoring proteins at the lipid-aqueous interface was shown to be a strong cholesterol binding motif which catalyzes the unfolding in the membrane. In the absence of cholesterol reduced insertion of the *β*-tongue is observed and the biased simulations reveal increased curvature and thinning of the POPC membrane consistent with the higher free energy associated with the transition in absence of cholesterol.

## Materials and Methods

The crystal structure of the ClyA monomer was taken from the PDB ID 1QOY^7^ (see Figure 1A), and the crystal structure of the protomer state was taken from the PDB ID 2WCD^8^ (see Figure 1B). The *β*-tongue (Figure 1C) and the helix-turn-helix motif (Figure 1D) were truncated from the monomer and the protomer state respectively. The amino acid sequence and the residues (170-207) are shown in Figure 1F. We worked with the crystal structure definition of the *β*-tongue, i.e., residue 177-203; however, the selected residues do not anchor to the membrane headgroups and lack the charge residue. Pore-forming toxins are known to have an evolution design in such a way that the charged residues reside at the edges of the transmembrane region, keeping the protein hinged to the lipid headgroup using electro-static interactions.^18, 19^ In the case of the ClyA, the *β*-tongue region lies from residue 177 to 203 in the monomer, which does not have any charged residues; however, a few additional residues on both sides of the peptide have several charged residues (Figure 1F). Previously, the protomer simulations of the ClyA embedded in the membrane have shown additional residue interaction with the membrane components.^16, 20^ Therefore, for our study, we consider residues from 170 to 207 of the *β*-tongue that were shown to interact with the membrane in the protomeric state, and their RKWY type of amino acid residues have electrostatic interaction with the lipid headgroup.

The initial structure of the bilayer was generated using the CHARMM-GUI membrane builder.^21^ Subsequently initial structures of *β*-tongue exposed to the membrane, *β*-tongue placed in the membrane, and the helix-turn-helix motif placed in the membrane were prepared. All-atom MD simulations for the membrane inserted and membrane exposed truncated *β*-tongue of monomer (Figure 1C), and the membrane inserted truncated helix-turn-helix motif (Figure 1D) of the protomer state were performed on a *1-palmitoyl-2-oleoyl-snglycero-3-phosphocholine* (POPC) membrane and a POPC membrane with 30% cholesterol (POPC:Chol) similar to the composition used in supported bilayer experiments in the single molecule study by Sathyanarayana et al. ^16^. All initial structures were solvated with TIP3P water and Na^+^ and Cl*^−^* ions were added to maintain the charge neutrality and 150 mM of salt physiological concentration. Initial solvated structures were energy minimized using the steepest-descent method with a maximum force of 1000 kJ mol*^−^*^1^ nm*^−^*^1^. The energy minimized structures were then gradually heated up to 310 K, followed by 10 ns of NVT simulation and 20 ns of NPT simulations with position restraints on protein, lipid, and cholesterol molecules. The final configurations from the equilibration simulation were used as the starting structures for the production runs. All simulations were performed in the NPT ensemble at 310 K using GROMACS-2018.6 with the force field parameters for the ClyA protein taken from AMBER99SB-ILDFN force field with phi corrections, and the force field parameters for lipid and cholesterol were taken from the compatible SLIPIDS force fields.^22–24^ This force-field combination has been shown to accurately reproduce the crystal structure of dodecameric pore complex in the membrane environment and compared well with other force-fields such as the CHARMM36 protein and lipid FFs as well as the AMBER class of force-fields^20, 25^ The semi-isotropic pressure control was achieved using the Parrinello-Rahman method with a time constant of 1 ps. The temperature was controlled using Nośe-Hoover chains with a time constant of 0.1 ps. The long-range electrostatic interactions were treated using the particle mesh Ewald (PME) method with a real space cut-off of 1.5 nm and van der Waals interactions computed with a cut-off of 1.5 nm. Three-dimensional periodic boundary conditions were applied to eliminate boundary effects. Hydrogen bonds were constrained using the LINCS constraint, which allowed a larger time step of 2 fs.^26^ The solvent, lipid, cholesterol, and protein molecules were coupled separately to a temperature bath at 310 K. Pressure was kept constant at 1 bar using the isothermal compressibilities, *K_xy_*= *K_z_*= 4.5 *×* 10*^−^*^5^ bar*^−^*^1^.^27^ Equilibration was monitored by evaluating the root mean squared deviation (RMSD) and secondary structure changes of the protein. Complete simulation details are given in Table S1.

### String method simulation details

Two stable endpoints of the string were carefully selected based on the multiple replicates of the equilibrium simulations of monomer and protomer states. For the POPC membrane, the two endpoints are the *β*-sheet (Figure S3A) and the helix-turn-helix motif (Figure S3B). This is in contrast to simulations with cholesterol where we observed spontaneous unfolding of the *β*-tongue and hence used the partially unfolded state as one of the endpoints of the string.^1^ The conformation phase space was defined using path collective variables (PCVs) (*S, Z*) originally developed by Branduardi et al. ^17^. The PCVs is defined for a system consist of *N* number of atoms with positions (**r**_1_, **r**_2_, …, **r***_N_*) *≡* **R** undergoing a conformational change. First, we consider a sequence of *n* reference structures, **R**^(1)^, **R**^(2)^, …, **R**^(*n*)^ of the molecule under consideration, connecting two endpoints of the conformational change. With the help of reference structures, PCVs are defined as,

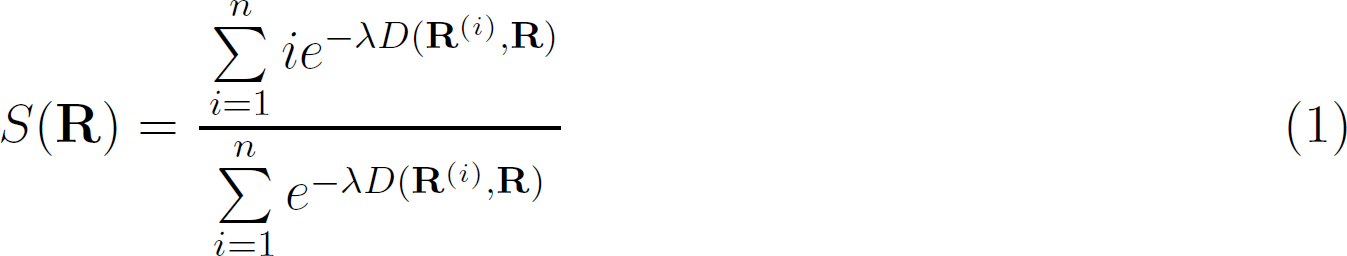

and

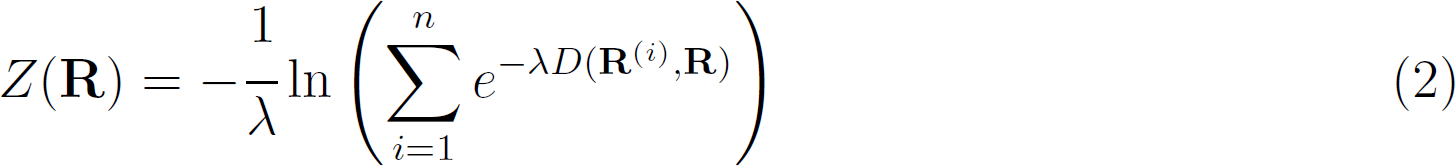

where *S* denotes the progress from one endpoint to the other along the reference path, *Z* is a measure of the variation in the perpendicular direction. In our work, the reference path consisting of 20 states for the PCVs was generated using linear interpolation between the atomic coordinates of the structures between the two end states. The *λ* values were 594.88 nm*^−^*^2^ for the POPC:Chol membrane and 422.00 nm*^−^*^2^ for the POPC membrane. These values are inversely proportional to the mean square displacement (MSD) between two consecutive states on the reference path.

Sampling from the unbiased simulations of the endpoints projected in (*S, Z*) space is shown in Figure S3C for the POPC membrane where the states are clearly distinguishable and indicate the presence of a free energy barrier. The string was defined by 50 points connecting initial and final states. To evolve the string, a force constant value of 100 kJ mol*^−^*^1^ was used in the *S* and *Z* variables. During string evolution, 0.5 ns of simulations were performed on each point in every iteration until the string converged. The convergence of the string was monitored using,

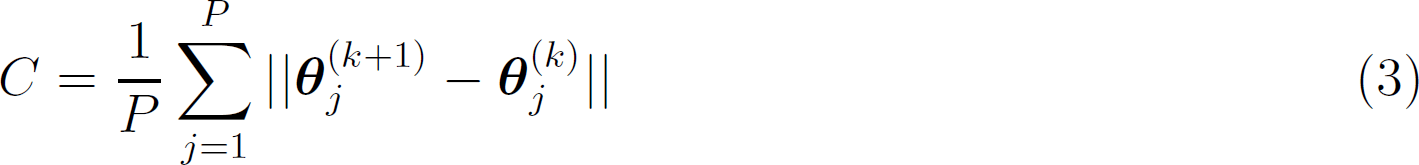

where, *θ_j_* represents the co-ordinates (*S_j_, Z_j_*) and *||·||* is the Euclidian norm. The superscript *k* is the iteration number, subscript *j* represents the point on the string, and *P* is the total number of points on the string. The string converged in 12 iterations for the POPC membranes (see Figure S3D). Additional 60 ns umbrella sampling simulations on each point of the converged path were performed to obtain a converged free energy landscape along the transition path. The overlapping of histograms for WHAM analysis was ensured by examining the regions with sampling counts greater than 0.1 million as illustrated in Figure S3F. The 1D free energy profile was reported on the arc length parameterized distance ‘P’ calculated from the monomer state to the protomer distance. The convergence of the free energy profile was monitored using the evolution of the free energy profile upon performing additional simulations at each point along the string. In Figure S3E, the free energy was found to be invariant beyond 45 ns of simulation.

### Simulation analysis

Gromacs in-built commands were used to compute root mean square deviation (RMSD), root mean square fluctuation (RMSF), and the radius of gyration (ROG). Time trajectories of secondary structure changes were monitored using Gromacs ‘do-dssp’ command and VMD timeline features with the STRIDE software. For the Chol and POPC atom-atom occupancy analysis with the protein, contact between two atoms was assumed to occur if the distance was less than a cutoff distance of 0.5 nm. ^28, 29^

The fractional occupancy *O_r_* for residue, *r* is defined using,

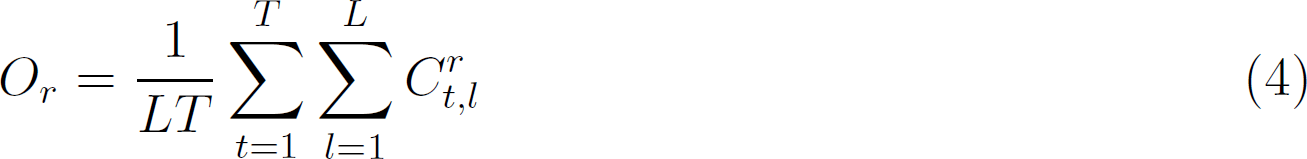

where *t* is the time index, *T* is the total simulation time, *l* is the index for number of lipid molecules, *L* is the total number of lipid molecules. 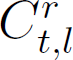 denotes the total number of contacts formed between *r^th^* residue and *l^th^* lipid molecule at time *t* and is evaluated using,

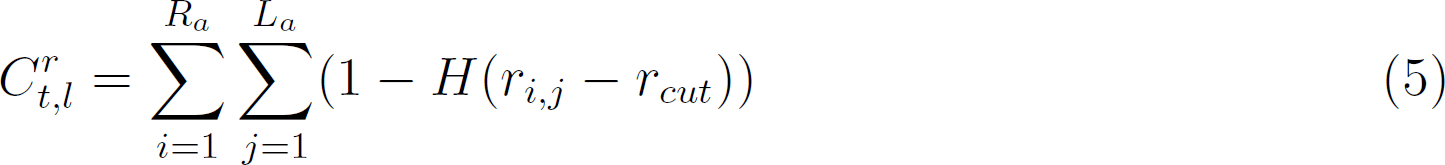

where *H*(*x − x_o_*) is the Heaviside step function whose values is 0 when *x < x_o_* and 1 when *x ≥ x_o_*.

The membrane curvature and the thickness were computed using the MDAnalysis Python package,^30^ where first, a reference group of atoms is selected to define the upper and lower membrane surfaces. We have chosen phosphorous atoms of upper and lower leaflets to define the respective surfaces. Every atom of the reference group is assigned a grid based on its *x* and *y* location. Once every grid is populated with the reference atoms, the *z* coordinates of the atoms are stored for each configuration. The average upper and lower surfaces were reported by averaging each grid value. The difference between the lower and upper leaflet surfaces was used to compute the membrane thickness, and the mean curvature was evaluated using the method proposed by Yesylevskyy and Ramseyer ^31^. The extent of penetration of the protein or residue in the membrane is carried out by analyzing the *z*-coordinate of the distance between the center of mass of the protein/residue and the center of mass of the membrane.

## Results and discussion

### Cholesterol promotes the unfolding of ***β***-tongue monomer state

We first carry out MD simulations to study the secondary structure changes of the *β*-tongue in POPC and POPC:Chol membranes (Figure 2). These simulations are required to determine the end states of the protein for the string method based free energy computations. Unlike in the case of POPC:Chol membranes^1^ where the protein partially unfolds and the *β*-sheet content is completely lost (Figure 2C), the structure of the *β*-tongue remained stable without appreciable loss of *β*-sheet content (Figure 2B) in the absence of cholesterol. However, in both cases, the *β*-tongue remained hinged to the lipid headgroups.

**Figure 2:**
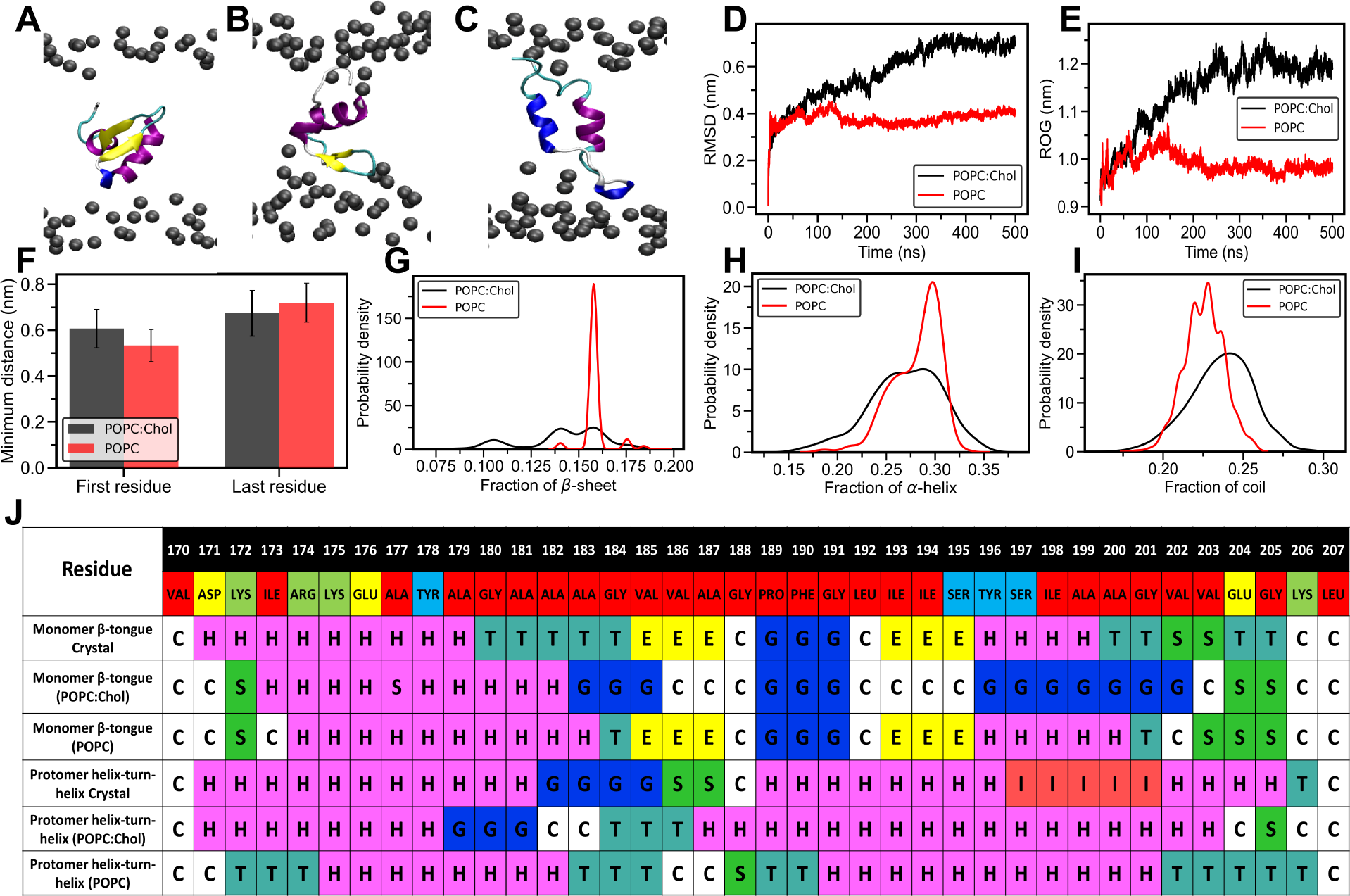
Monomer *β*-tongue placed in the membrane. (A) Initial state of the *β*-tongue placed in the membrane. The final structure of the *β*-tongue at the end of the 500 ns simulation for the (B) POPC membrane and (C) POPC:Chol membrane. (D) Average RMSD and (E) average ROG comparison for POPC:Chol and POPC membrane averaged over three independent 500 ns simulations. (F) The minimum distance between the C-*α* atom of the first (170) and last residue (207) to the lipid headgroups. Fraction of secondary structure content for (G) *β*-sheet, (H) *α*-helix, and (I) coil structure. (J) Residue-wise (Red: Non-polar, Blue: Polar, Green: positively charged, Yellow: Negatively charged) secondary structure in crystal structures and different membrane environments. Color and string codes are as follows; pink: *α*-helix (H), teal: turn (T), yellow: *β*-sheet (E), blue: 3_10_-helix (G), red: *π*-helix (I), green: bend (S), white: coil (C). RMSD and ROG data for POPC:Chol adapted from Kulshrestha et al. ^1^, Copyright 2023, American Chemical Society.

The average behavior of the RMSD from three independent MD simulations (Figure 2D) shows a higher RMSD of around 0.7 nm for the POPC:Chol membrane and 0.4 nm for the POPC membrane after 300 ns of the simulation. The radius of gyration (ROG) (Figure 2E) also suggests that the protein remains more compact in the POPC membrane with a lower value of ROG. In order to quantify the hinged state of the protein as illustrated in Figures 2B and 2C, we evaluated the averaged minimum distance between the C*α* atom of the first (VAL170) and last (LEU207) residues of the *β*-tongue with the phosphorous atoms of the lipids (Figure 2F). The data illustrates that both the terminal residues are in close proximity to the lipid head groups independent of the presence of cholesterol. Once hinged, they remain attached to the headgroup with a distance of around 0.5-0.6 nm for the first residue (VAL) and 0.6-0.7 nm for the last residue (LEU). Both these are adjacent to charged residues ASP171, LYS172, and LYS206, which can form electrostatic interactions with the lipid headgroups to stabilize these hinged configurations. These electrostatic interactions play a broader role across membrane proteins and residues such as ARG, LYS, TRP, and TYR have been shown to stabilize protein interactions in the headgroup region at the protein-membrane interfaces in pores formed by ClyA and *α*-hemolysin.^19^

The fraction of secondary structure content (Figure 2G - 2I) also indicates a distinct unfolding of the *β*-sheet in the presence of cholesterol where increased conversion into a coil structure is observed. In contrast, the secondary structure content is more rigid in the absence of cholesterol and this tendency is the greatest in the case of the *β*-sheet (Figure 2G). The representative differences in the secondary structure content for the *β*-tongue residues are summarized in Figure 2J for configurations taken at the end of a single 0.5*µ*s simulation. For the monomer *β*-tongue in the POPC:Chol membrane, we observe an increase in 3_10_ helices (G) and coils (C) with a complete absence of *β*-sheets (E) when compared with the crystal structure. In contrast the *β*-tongue in POPC retains similar secondary structure to the crystal state. In the POPC membrane the resistance of the *β*-tongue to unfold suggests the presence of a barrier to form the partially unfolded intermediate state and hence to refold to the helix-turn-helix motif of the protomer state. Since we are interested in capturing the mechanism for the conformational change, we select the *β*-tongue state (Figure 2B) as one endpoint of the string for the free energy analysis in the POPC membrane. We note that in earlier free energy computations in the POPC:Chol membrane, a partially unfolded state was used as one endpoint for the free energy analysis since this state was formed spontaneously in the MD simulations.

To define the second end state of the string for the free energy analysis, we simulated the helix-turn-helix motif (residues 170-207) of the protomer, which is the stable conformation of the membrane inserted pore state, placed in the POPC:Chol and POPC membranes (Figure Figure S1). The protomer state in both membranes remained stable and was found to be hinged to the lipid headgroups as illustrated in Figure S1F. These simulations indicate that the helix-turn-helix motifs are stable conformations that could be used as end points for the string method analysis. The secondary structure analysis (Figure 2J) also revealed that the the helix-turn-helix motifs for the protomer states are preserved in both membranes.

We next apply the string method to capture the transition from the *β*-sheet state (Figure 2C) to a protomer state (Figure S1B) in the POPC membrane and use a similar method based on path collective variables (PCVs)^17^ as outlined in our previous work.^1^

### Unfolding of ***β***-tongue is on-pathway intermediate of the conformational change

In Figures 3A and B, we illustrate the converged paths obtained from the string method analysis for the POPC and POPC:Chol membranes respectively. In order to facilitate a comparison between the two membranes, we have adapted the free energy data from our previous work and computed several additional properties to quantify the changes along the string for the POPC:Chol membrane.^1^ The converged strings to the transition path for both cases remain below 0.2 nm^2^ in *Z*, indicating that the transition path does not deviate significantly from the reference path defined at *Z* = 0. In case of the POPC:Chol membrane, the initial state at *S* = 1 corresponds to the partially unfolded state, and *S* = 20 corresponds to the final protomer state. For the POPC membrane *S* = 1 corresponds to the *β*-sheet state of the monomer in the POPC membrane as determined from unrestrained MD simulations discussed earlier. Representative snapshots of the structures on the converged path are shown in Figure 3. These structures were generated using additional short biased MD simulations for 3 ns using harmonic potentials for *S* and *Z* with 1000 kJ mol*^−^*^1^ nm*^−^*^1^ spring constants. For all points sampled along the path, we observe that N (green) and C (red) termini remain hinged to the lipid headgroups for both membranes. For the POPC membrane we also observe large deviations in the phospholipid headgroups from a planar configuration. The influence of the protein on the topological changes to the membrane surface is quantified and discussed later in the text.

**Figure 3:**
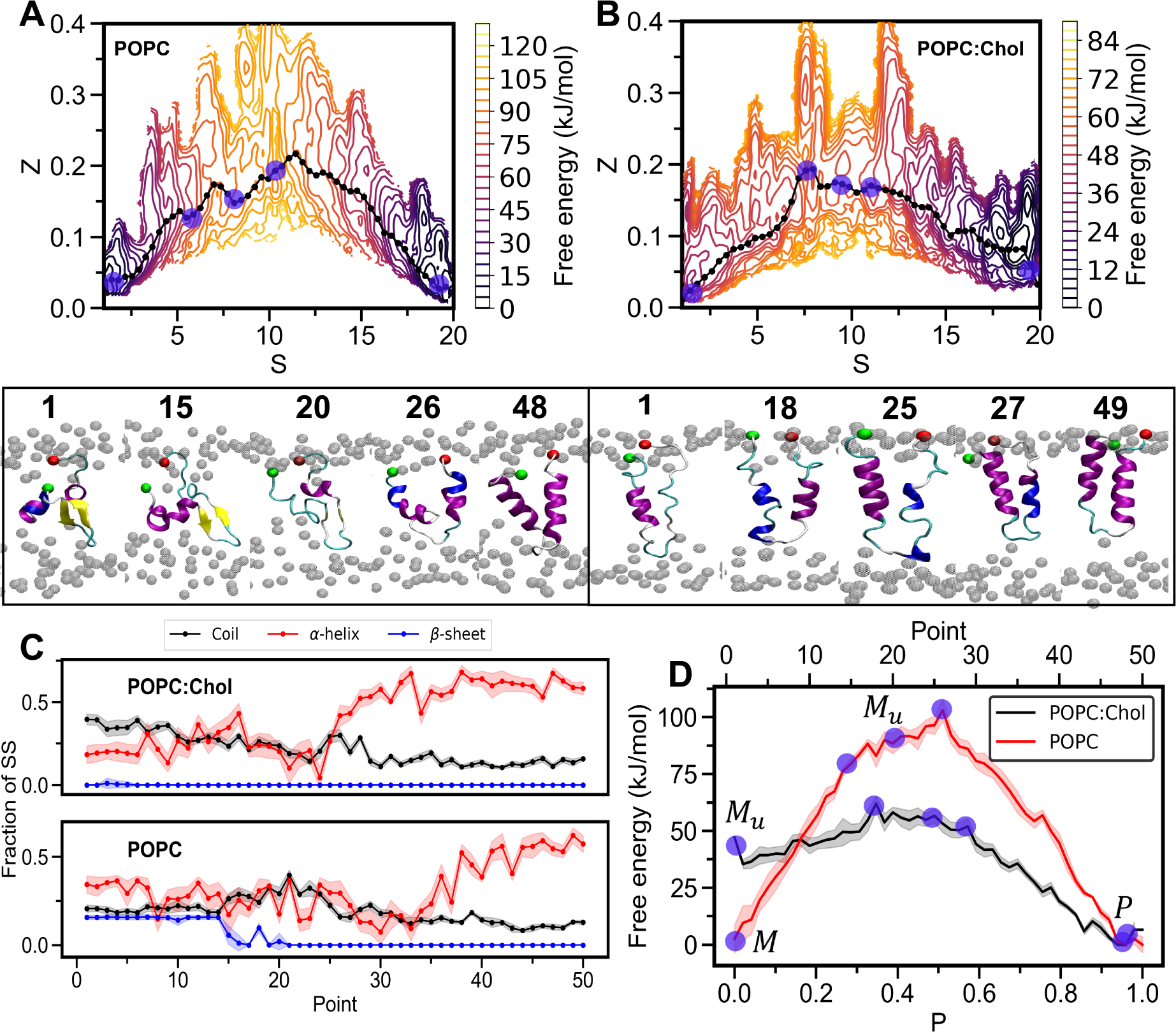
Transition path of ClyA *β*-tongue and the free energy profile. The final converged string (black color) and the free energy surface contour plot for the case of (A) the transition from the *β*-tongue to the protomer state in POPC membrane, and (B) the transition from the unfolded state to the protomer state in the membrane with cholesterol. A few representative snapshots of the structure on the path are shown below, where the green sphere represents the N-terminus, and the red sphere represents the C-terminus. The values of (*S, Z*) for the corresponding point on the string are given in Table S2 and Table S3 for POPC:Chol and POPC membrane, respectively. (C) Fraction of secondary structure changes on the transition path. (D) 1D free energy profile along the parameterized distance ‘*P* ’ calculated from partially unfolded state to the protomer state for POPC:Chol membrane, and *β*-sheet to protomer state for POPC membrane. Transparent blue circles in A, B and D are the location of the different snapshots. Free energy data in A and D for POPC:Chol adapted from Kulshrestha et al. ^1^, Copyright 2023, American Chemical Society.

An analysis of the secondary structure changes on the converged path along the string is shown in Figure 3C with the corresponding free energy profiles in Figure 3D. A detailed stride analysis of the secondary structure evolution for the final configurations on the converged string is given in the SI Figure S4. In the POPC membrane, the *β*-sheet content is lost between points 1-20 on the string, and an increase in the formation of turns is observed in this region. Reduced *β*-sheet content is observed in the corresponding snapshot shown at point 15 and the free energy rises to about 80 kJ/mol at this point. Subsequently, we observe fluctuations in the helical content indicative of unfolding (see snapshot point 27) and a further rise in the free energy till it reaches a maximum at point 26. Finally, the C-terminus strand gains helicity where it fluctuates between 3_10_-helix and *α*-helix (SI Figure S4) until it completely converts into *α*-helix. At later points we observe a refolding to form the helix-turn-helix motif of the protomer and this is accompanied by a decrease in the free energy to the protomer state. Thus the primary barrier for the transition in the absence of cholesterol lies in the unfolding of the *β*-sheet. Once unfolding is maximized, the refolding transition is a downhill in the free energy profile. In case of the POPC:Chol membrane, secondary structure data along the points confirm the presence of a partially unfolding transition along the string, however a weak barrier in the free energy which forms between points 20-30 is only *∼* 10 kJ mol*^−^*^1^ reflecting the strong role played by cholesterol in spontaneously unfolding the *β*-sheet. Refolding takes place beyond point 24 accompanied by a decrease in the free energy.

### Cholesterol stabilizes the protomer state and enhances the kinetics of the conformational change

Figure 3D, illustrates the differences in the free energy landscape for the POPC and POPC:Chol membranes. We propose the transition from monomer (M) to protomer (P) follows a two step process passing through a partially unfolded monomer intermediate M*_u_*.

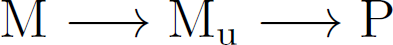

Since the partially unfolded state M_u_ depleted of *β*-sheet content in the POPC:Chol membrane was found to be a stable intermediate state, we ascribe the point on the string with the zero *β*-sheet content to define the corresponding intermediate M_u_ for the POPC membrane, which occurs at point 20 on the string (Figure 3C) lying below the maximum in the free energy.

Tabulated values of the free energy changes are given in Table 1 where the barrier to form M*_u_* from the M state (ΔG_1_ = G*_M__u_* - G*_M_*) in the POPC membrane is 85.55 kJ mol*^−^*^1^. Since M*_u_* forms spontaneously at room temperature for the POPC:Chol membrane, we assign the barrier to be of order k*_B_*T. The free energy difference (ΔG_2_ = G*_P_* - G*_M__u_*) to transform from M*_u_* to the protomer state is about three times lower for the POPC:Chol membrane. Upon closer inspection, the free energy profiles reveal that the protein from the M*_u_* state must overcome a small barrier during the transformation to the helix-turn-helix protomer state.

**Table 1:** Free energy data. Free energy differences ΔG are reported for transitions between the monomer ‘M’, protomer ‘P’, and partially unfolded monomer ‘M_u_’ states. See text for definition of different quantities.

| Membrane | $\Delta G_1$ kJ/mol | $\Delta G_2$ kJ/mol | $\Delta G_3$ kJ/mol |
| --- | --- | --- | --- |
| POPC | $85.55 \pm 10.9$ | $-88.44 \pm 7.9$ | $14.66 \pm 7.9$ |
| POPC:Chol | $\sim k_B T$ | $-35.43 \pm 2.49$ | $26.56 \pm 5.30$ |

This barrier is associated with an additional loss of secondary structure due to a decrease in *α*-helical content and an increase in turns and 3_1_0 helices (Figure 3D and Figure S4). We evaluate the corresponding free energy change as the difference between the transition state, i.e., the maximum (point 26 for POPC and 18 for POPC:Chol) and the free energy of the M*_u_* state (ΔG_3_ = G*_max_* - G*_M__u_*). The free energy barrier for this secondary transition is about a factor of two greater for the POPC:Chol membrane. Once this barrier is surmounted, refolding to the protomer state is a downhill process. Therefore, for the POPC membrane, the largest barrier occurs during the unfolding stage of the monomer in the membrane (Δ G_1_). These barriers that we observe are related to the kinetics of the transitions that occur when the protein binds to the membrane. Our study with the *β*-tongue motif sheds light on only one part of the conformational changes. The complete transition also involves the insertion of the N-terminal helix into the membrane (Figure 1). We draw qualitative connections with experiments carried out with ClyA with the present free energy analysis. The kinetics of pore formation vary widely depending on the specific membrane platform and assay used.^16^ In dye leakage experiments with small unilamellar vesicles, the leakage kinetics increased by a factor of three in the presence of cholesterol. Interestingly in POPC:Chol vesicles, the kinetics was invariant for cholesterol content ranging from 30-50% indicating the presence of a threshold cholesterol concentration for robust pore formation. In contrast, turbidity assay experiments with ClyA showed a 100 fold increase in rupture activity when cholesterol was present. These differences result in part from the large curvature present in the vesicle leakage experiments, which retard the pore formation kinetics. Despite these variations, our free energy computations indicate the lowered barrier in the presence of cholesterol consistent with the enhanced leakage kinetics observed in the experiments.^16, 32, 33^

### Tyrosine is essential for the unfolding transition with cholesterol

In order to obtain molecular insights into the role of cholesterol, we analyzed the structures and interactions along the transition path using several different metrics. In Figure 4A, the average cholesterol occupancy computed from the last 50 ns of the biased trajectory on the converged path indicates that Tyr178 has the greatest occupancy, followed by Arg174 in the initial points along the path. However cholesterol interactions have a broader distribution across the residues along the path and in the protomer state several residues show strong cholesterol binding sites. These sites correspond to cholesterol hot spots observed by us earlier in simulations of the dodecameric ClyA pore complex in a POPC:Chol membrane ^16^ where cholesterol was found to bind between pockets formed by adjacent membrane inserted *β*-tongue motifs. In a recent mutagenesis and thermal unfolding study, we have shown a near complete abrogation of lytic activity of the mutant Tyr178Phe, in erythryocyte turbidity and the vesicle leakage assays. Furthermore, this mutant was resistant to thermal unfolding in membranes containing cholesterol,^1^ revealing that conformational flexibility was an important factor in assisting secondary structure changes involved during ClyA pore formation. Since the Tyr178 was found to be a key residue (Figure 4A), we analyzed the *β*-tongue, Tyr178Phe mutant trajectories from our earlier study^1^ to compare the cholesterol occupancies (Figure 4B). Tyr178Phe showed significantly reduced cholesterol occupancy at residues 174 and 178, however increased occupancy occurred at other residues of the *β*-tongue. For comparison we also computed the cholesterol occupancy for the wild type *β*-tongue with and without restraints to discern the differences between a folded and unfolded state respectively - noting that the wild type spontaneously unfolds in the presence of cholesterol. Tyr178 is clearly a cholesterol binding hot spot when unfolding is prevented using a restraint. A snapshot from the simulation (see inset in Figure 4B) shows the binding of the *α* face of cholesterol with tyrosine. In the absence of restraints where the wild type partially unfolds, cholesterol occupancy at Tyr178 is drastically reduced (Figure 4B). From this analysis, we hypothesize that cholesterol interaction with Tyr178 is critical to catalyzing the spontaneous unfolding of the *β*-tongue, and stabilizing the partially unfolded intermediate. Using replica exchange molecular dynamics, Miller et al. ^34^ has shown that the presence of tyrosine in the cholesterol recognition and consensus motif (CRAC) motif of leukotoxin facilitates unfolding of the peptide, suggesting that the interaction with tyrosine, a necessary residue for the definition of the CRAC motif, is essential for this transition. We point out that the tyrosine present in the *β*-sheet motif is not part of a CRAC motif implicated in cholesterol-protein interactions.^35^

**Figure 4:**
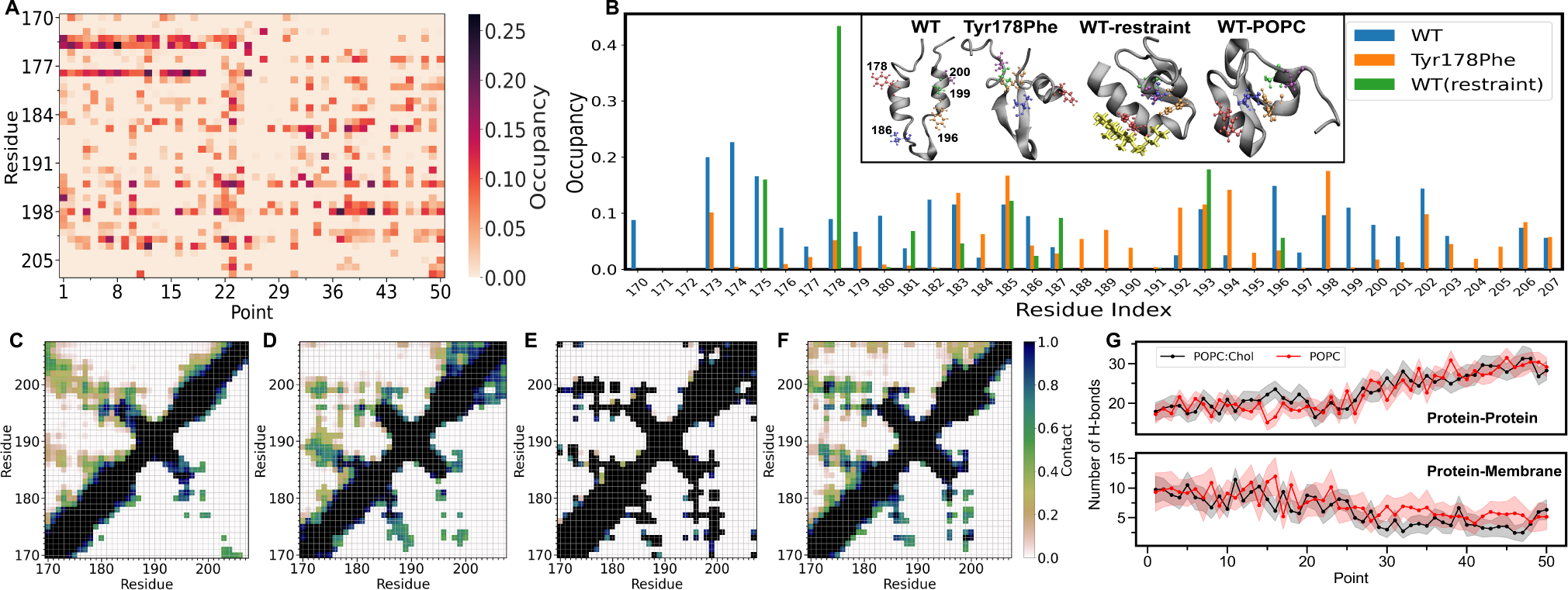
Conformational changes of the ClyA *β*-tongue. (A) Average cholesterol occupancy comparison for each residue of monomer *β*-sheet in POPC:Chol membrane on the transition path. (B) Residue-wise average cholesterol occupancy for wild type (WT) and Tyr178Phe mutant. Inset figures represents the snapshots taken at the end of the 500 ns MD simulations, where the residue 178, 186, 196, 199, and 200 are colored red, blue, orange, green, and purple respectively. Cholesterol molecule interacting with restraint WT is shown in yellow color. Contact map of protein for (C) WT, (D) Y178F mutant, (E) WT restraint, and (F) WT in POPC membrane. Lower triangular regions of contact maps are plotted only for contact values greater 0.5 to identify to strong contacts. (G) The number of hydrogen bonds between protein-protein and protein-membrane, where the membrane is composed of POPC and cholesterol molecules.

The contact maps of protein based on the distance between C*_α_* atoms ¡ 1 nm (Figures 4C to 4F) reveals the role of cholesterol in the WT protein during *β*-tongue unfolding. The lower triangular part of the contact maps only depicts strong contacts defined as those that persist for greater than 50% of the simulation time. The contact map of the partially unfolded WT shows a loss of all strong contacts (Figure 4C) in comparison to the WT protein restrained to prevent unfolding (Figure 4E). In the case of the restrained WT, residue 178 makes strong contacts with residues 186, 196, 199, and 200 (Figure 4E). WT ClyA in the POPC membrane (Figure 4F) maintains largely the same strong contacts (between 178 residue and residues 186, 199, and 200) suggesting that cholesterol is involved in the unfolding observed in restraint-free simulations. Similarly, Tyr178Phe mutant displays a comparable contact map in the vicnity of residue 178 residue with strong contacts with residue 186 (Figure 4D) providing additional evidence that Tyr178 is indeed necessary for cholesterol mediated unfolding. Overall, our contact map analysis along with the occupancy analysis suggest that the Tyr178 has a strong affinity to interact with the *α* face of the cholesterol, and its interaction with cholesterol disrupts the internal contacts within the *β*-tongue motif leading to the unfolded state with reduced contacts. Once the protein is unfolded other residues are accessible to cholesterol but none reveal a strong propensity to compete out the others (Figure 4A). Consistent with the idea that the major role of cholesterol is to induce unfolding, the occupancy at residues 174/178 disappear at point 19 (Figure 4A) concomitant with loss of *α*-helix and *β*-sheet content (Figure 3C).

The variation in the number of hydrogen bonds (H-bonds) between protein atoms, and protein and membrane atoms are shown in Figure 4G. The number of protein-protein H-bonds remains *∼* 20 till point 25 along the string for both the POPC and POPC:Chol membranes, with a subsequent increase associated with refolding to the protomer state. This refolding is found to occur around point 25 on the string for the POPC:Chol membrane and a little later for the POPC membrane (Figure 3). However, we were unable to discern the differences in the H-bond data between the two membranes. Concomitant with the increase in H-bonds for the protein-protein interactions, we observe a decrease in protein-membrane H-bonds.

### Cholesterol facilitates the ***β***-tongue insertion

The study has thus far focussed on the conformational changes that occurs when the *β*-tongue is inserted in the membrane. However, it is also of interest to study the extent of secondary structure changes when the monomer first binds to the membrane. Since membrane binding is primarily mediated by the *β*-tongue^8, 20^ we carried out MD simulations with the *β*-tongue initially placed in the vicinity of the phospholipid headgroups for both POPC:Chol and POPC membranes (Figure 5A). In Figure 5B, the fraction of secondary structure analysis reveals a distinct loss of *β*-sheet content in the presence of cholesterol in the membrane, consistent with the behavior observed when the *β*-tongue was inserted the membrane, reiterating that unfolding is on-pathway to the conformational change accompanying the monomer to protomer transition. The final snapshots in Figure 5C illustrate an increased unfolding of the protein in the POPC:Chol membrane along with increased membrane penetration. Furthermore, residue-wise depth analysis in Figure 5D indicates that the residues toward the N-terminus lie deeper in the POPC:Chol membrane when compared with the POPC membrane. It is important to note that the N-terminus side of the monomer consists of many charged residues (Figure 1F), which have enhanced cholesterol binding (Figure 4A).

**Figure 5:**
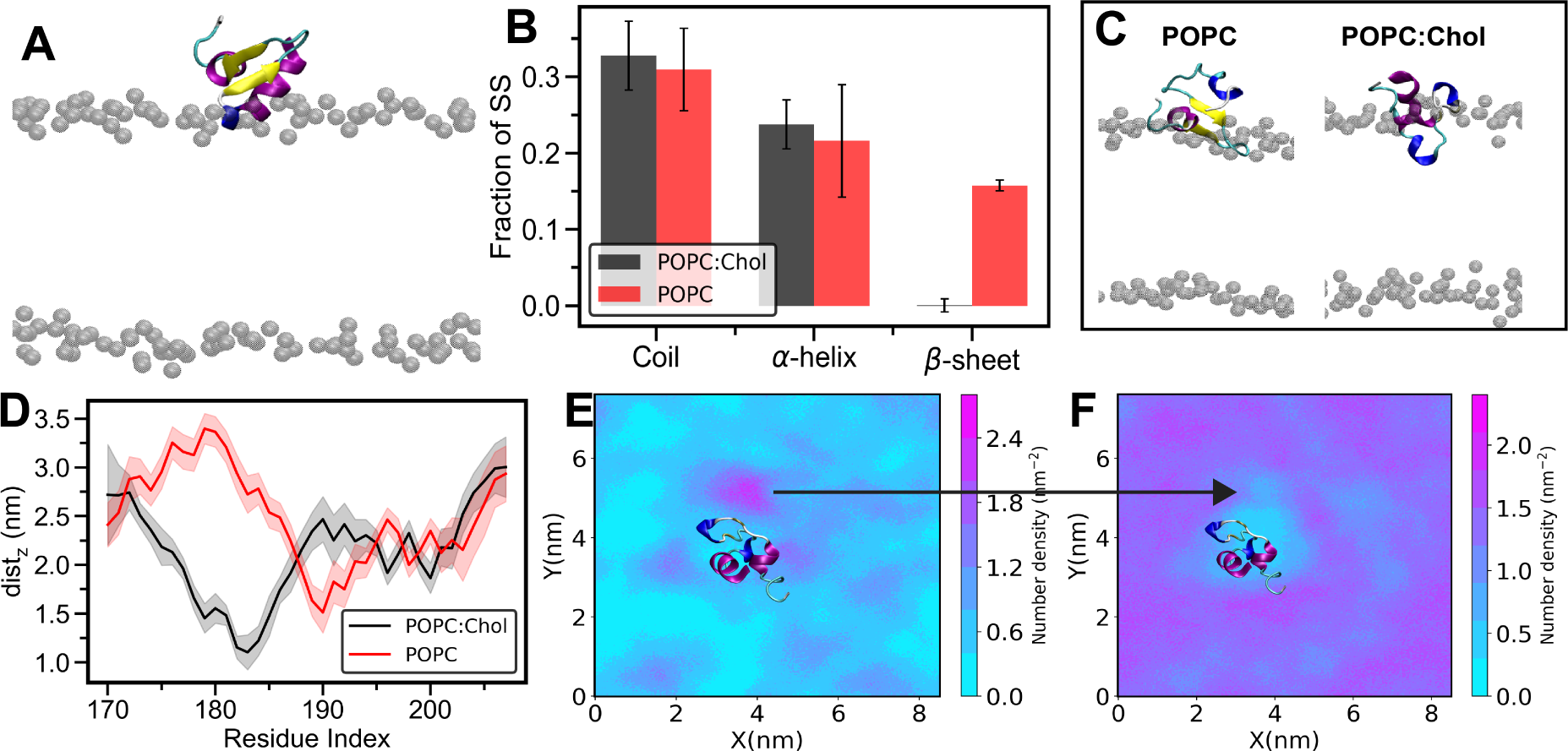
*β*-tongue exposed to the membrane. (A) Initial state of the *β*-tongue exposed to the POPC:Chol and POPC membrane, with only the location of the headgroups illustrated for clarity. (B) Fraction of secondary structure content. (C) Final snapshots of the protein in the membrane at the end of one microsecond-long simulation. (D) residue-wise depth analysis with the *z*-coordinate of the distance between the center of mass of a residue and the center of mass of the membrane. Number density map for (E) cholesterol molecules and (F) POPC molecules in the upper leaflet of the POPC:Chol membrane system, the black arrow indicates the accumulation of the cholesterol near the protein and displacement of the POPC molecules.

We have also simulated the *β*-sheet in solution to assess the protein stability in the absence cholesterol. In Figure S2, the RMSD analysis indicates that the protein is well equilibrated within 200 ns of simulations, and the *β*-sheet remains stable as illustrated in Figure S2B. Interestingly, the helical region of the protein first unfolds and then converts into a *β*-sheet resulting in the increase in the *β*-sheet secondary structure content. This observation further supports our view that the innate unfolding of the *β*-sheet in the POPC:Chol membrane is catalyzed by the presence of cholesterol.

We analyze the redistribution of cholesterol and phospholipids around the protein using the 2D number density maps illustrated in Figures 5E and 5F respectively. This data is computed from MD simulations where the *β*-tongue is placed at the membrane interface (Figure 5E) and averaged over the last 500 ns of the 1 *µ*s restraint free simulations. The number density reveals the enhanced density of cholesterol molecules around the protein suggesting that preferential binding of cholesterol occurs prior to the complete insertion of the *β*-tongue in the membrane playing a key role in facilitating the *β*-tongue insertion as observed in the POPC:Chol membrane. We point out that MD simulations of the complete ClyA monomer placed at the phospholipid interface^1^ with the *β*-tongue in close proximity to the headgroups, also showed the partial unfolding and insertion of the *β*-tongue over a 1 *µ*s simulation in the POPC-Chol membrane indicating that these trends are generic and consistent with the trends observed with the isolated *β*-tongue motif reported here.

### Membrane modulation with cholesterol

Using the mean position of the phosphate atoms for the upper and lower leaflets (Figure 6) we examined the influence of different conformations of the *β*-tongue motif on membrane thickness variations and ensuing deformations. The surface representations of the local membrane thickness are illustrated in Figures 6A-6D for the initial monomer and final protomer states of the protein used for the string method computations. Analysis of the curvature for different points along the string reveals similar differences between the two membranes. Cholesterol is known to increase the thickness of a POPC membrane^36^ and we observe a similar increase in the thickness in our simulations. In the presence of the *β*-tongue significant differences in the local membrane thickness variations are observed between the POPC and POPC:Chol membranes. The average thickness of the membrane with the protein reduced from 40.42 *±* 0.53 nm to 36.57 *±* 0.55 nm in the absence of the cholesterol. In the POPC membrane, the presence of the protein results in large curvature and membrane thickness modulations across both leaflets of the membrane. The upper leaflet shows a large positive mean curvature (Figure 6E) with a corresponding negative mean curvature in the lower leaflet. Additionally, the local curvature and height variations are greatest in the vicinity of the protein (Figures 6E) and occur in both the monomer and protomer states (Figures 6A-6D). In contrast, the mem-brane thickness and curvature variations in the POPC:Chol membranes are less, however we observe smaller topological variations in the upper leaflet where the N and C termini are anchored. The increased free energy penalty due to larger induced curvature^37^ in the membrane for the POPC bilayer is consistent with the higher free energy for the transition from the monomer to the protomer in the absence of cholesterol. Thus the barrier for unfolding of the protein is also influenced by the extent of membrane deformation exacerbated in the absence of cholesterol. Proteins, especially curvature-inducing proteins, have been found to be remodel the membrane followed by preferential accumulation of curvature-sensing proteins on the curved surface.^38^

**Figure 6:**
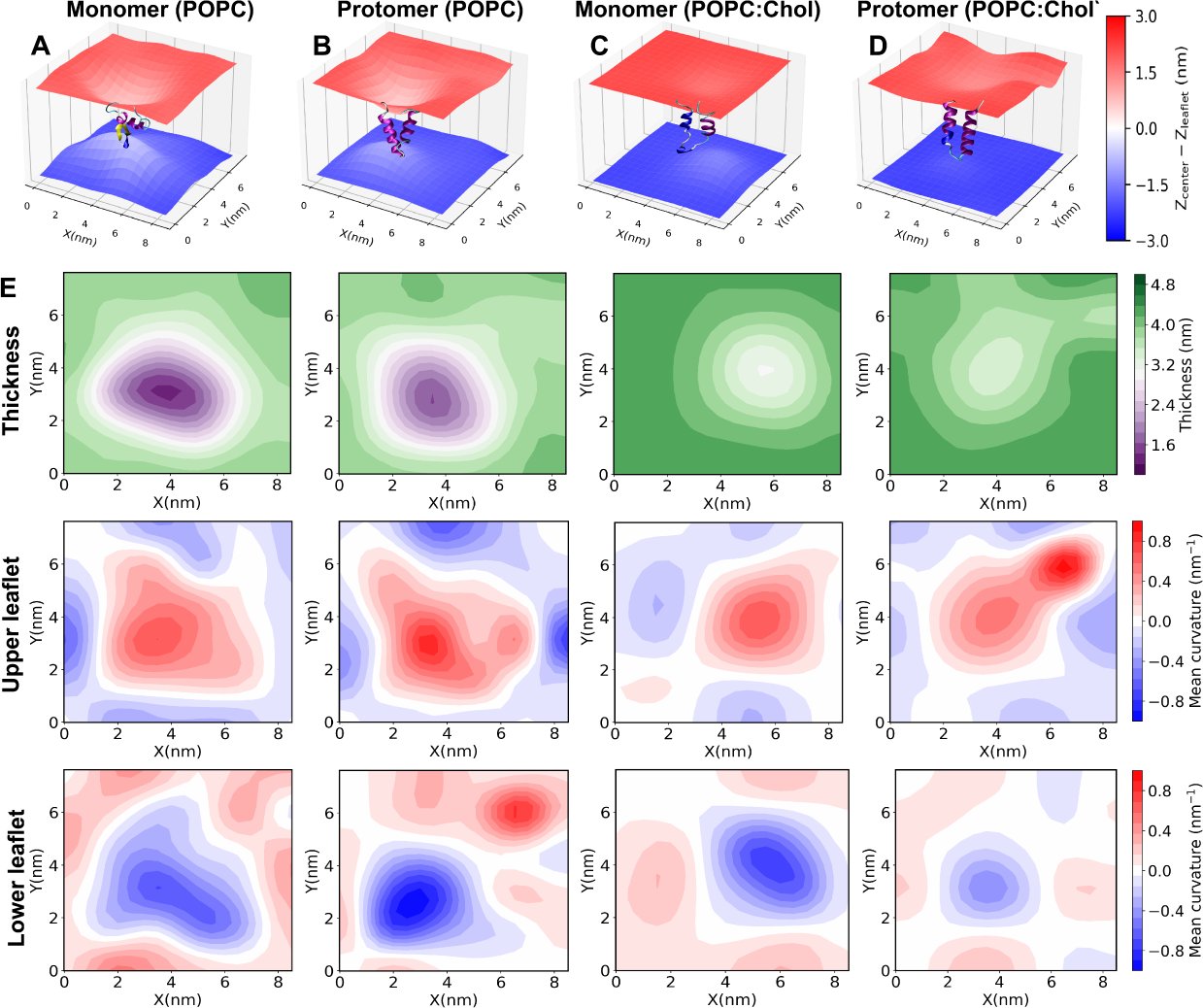
Membrane modulation. Upper extracellular and lower cytosolic leaflet mean surface representations for (A) monomer, (B) protomer placed in POPC membrane, and (C) monomer, (D) protomer placed in POPC:Chol membrane. *z_center_* is the average *z*-coordinate of the center of the membrane, and the *z_leaflet_* is the average surface coordinate of a leaflet. Red denotes the upper leaflet and blue denotes the lower leaflet surfaces. (E) Thickness and the upper and lower leaflet mean curvatures for POPC and POPC:Chol membranes.

## Conclusion

The ClyA pore forming toxin undergoes one of the largest conformational changes during membrane binding and pore formation. The predominantly hydrophobic *β*-tongue motif inserts into the membrane and undergoes a secondary structure transition to a helix-turn-helix motif. Kinetics of ClyA pore formation is known to be facilited by cholesterol, however the molecular underpinnings for this transition are poorly understood. Here we report for the first time an extensive free energy analysis using the string method of the folding that occurs in the *β*-tongue motif of ClyA in phospholipid membranes to unravel the specific role played by cholesterol in this transition. Our simulations reveal the presence of a large barrier for the *β*-tongue to unfold in the absence of cholesterol confirming that unfolding of this transmembrane pore motif is a critical step in the pore formation pathway of ClyA. Combined with our earlier work where we reported the free energy change for the *β*-tongue in the presence of cholesterol^1^ we posit that cholesterol catalyzes and stabilizes the unfolded intermediate which subsequently refolds to complete the conformational change. Refolding to form the helix-turn-helix motif in both POPC and POPC:Chol membranes is a favourable downhill transition on the free energy path. Analysis of the secondary structure content along the path reveals that the *β* sheet structure is resistant to unfolding in the absence of cholesterol and spontaneously unfolds in the presence of cholesterol. More specifically, initial binding of cholesterol to the residue Tyr178 in the *β*-tongue motif appears to trigger the unfolding in the membrane. This is consistent with the observation that a single point mutation Tyr178Phe was found to severely abrogate ClyA pore formation activity and MD simulations also showed the resistance to unfold in a POPC:Chol membrane^1^ for this mutant. Therefore cholesterol is able to compete for interactions of with Tyr178 to interfere with its role in stabilizing the folded *β*-tongue but also fine-tuned such that it no longer interferes in refolding after the structure has unfolded. The larger barrier to unfold in the absence of cholesterol is accompanied by membrane topological deformations that result in increased membrane curvature and thinning.

It is well established that cholesterol-dependent cytolysin (CDCs)^12^ use cholesterol molecules as their receptor. However, ClyA an *α*-toxin that does not belong to the CDC family of toxins shows a higher activity in the presence of cholesterol. In an earlier MD simulation cholesterol was found to bind to pockets formed by adjacent *β*-tongue motifs thereby stabilizing the dodecameric ClyA pore complex.^16^ In light of the findings in this work we are able to provide a more complete story of the complex role played by cholesterol in the pore formation pathway for ClyA. In addition to stabilizing the pore complex, cholesterol facilitates the initial unfolding of the *β*-tongue thereby lowering the barrier for the transition to the helix-turn-helix motif of the pore state. We posit that this is the primary driver for the enhanced pore formation and lysis kinetics observed in the presence of cholesterol.^16^ We finally comment on the initial binding step that precedes membrane assisted unfolding and refolding. Although we have not carried out free energy computations of the initial *β*-tongue binding, our restraint free MD simulations suggest that cholesterol triggers the loss of secondary structure during the initial binding stage, assisting in greater membrane penetration. In addition to the specific problem of secondary structure transformation for the ClyA pore forming toxin, our findings suggest that cholesterol a necessary ingredient for several active processes in the plasma membrane, ^39^ plays a critical role in secondary structure transitions and folding of proteins in a membrane environment. Our study illustrates that path based methods with appropriate collective variables provides an efficient and tractable means of computing the free energy associated with these transitions. The method can potentially be extended to study a wide class of membrane-protein folding events.

## Author Contributions

**Avijeet Kulshrestha:** Molecular dynamics simulations, post-processing, data curation, interpretation, writing original draft, review, and editing. **Sudeep Punnathanam, Rahul Roy, K Ganapathy Ayappa:** Conceptualization, interpretation of results, manuscript writing, review, and editing.

## Acknowledgments

We acknowledge the Supercomputer Education and Research Center (SERC) computing facility, Thematic Unit of Excellence on Computational Materials Science (TUE-CMS) a Department of Science and Technology (DST) supported computing facility at the Indian Institute of Science Bangalore, and the National Supercomputing Mission, India for funding used in this work.

## Supplementary Material

Supplementary information can be found in a separate document.

**Table S1** ClyA *β*-tongue complete Simulation details.

**Table S2** Converged string location for POPC:Chol membrane.

**Table S3** Converged string location for POPC membrane.

**Figure S1** Simulations of the helix-turn-helix motif of the protomer in membrane.

**Figure S2** Solvated *β*-tongue simulation.

**Figure S3** String method detail For the membrane without cholesterol (POPC).

**Figure S4** Secondary structure change on the path comparison between POPC and POPC:Chol membrane.

## References

[1] Kulshrestha, A.; Maurya, S.; Gupta, T.; Roy, R.; Punnathanam, S.; Ayappa, K. G. Conformational flexibility is a key determinant of the lytic activity of the pore-forming protein, Cytolysin A. Journal of Physical Chemistry B 2023, 127, 69–84.

[2] Blazek, A. D.; Paleo, B. J.; Weisleder, N. Plasma membrane repair: a central process for maintaining cellular homeostasis. Physiology 2015, 30, 438–448.

[3] Albers, R. W. W. Basic Neurochemistry; Elsevier, 2012; pp 26–39.

[4] Cymer, F.; Von Heijne, G.; White, S. H. Mechanisms of integral membrane protein insertion and folding. Journal of Molecular Biology 2015, 427, 999–1022.

[5] Dal Peraro, M.; Van Der Goot, F. G. Pore-forming toxins: ancient, but never really out of fashion. Nature Reviews Microbiology 2016, 14, 77–92.

[6] Bischofberger, M.; Iacovache, I.; Van Der Goot, F. G. Pathogenic pore-forming proteins: function and host response. Cell Host & Microbe 2012, 12, 266–275.

[7] Wallace, A. J.; Stillman, T. J.; Atkins, A.; Jamieson, S. J.; Bullough, P. A.; Green, J.; Artymiuk, P. J. E. coli hemolysin E (HlyE, ClyA, SheA): X-ray crystal structure of the toxin and observation of membrane pores by electron microscopy. Cell 2000, 100, 265–276.

[8] Mueller, M.; Grauschopf, U.; Maier, T.; Glockshuber, R.; Ban, N. The structure of a cytolytic *α*-helical toxin pore reveals its assembly mechanism. Nature 2009, 459, 726–730.

[9] Giri Rao, V. H.; Desikan, R.; Ayappa, K. G.; Gosavi, S. Capturing the membrane-triggered conformational transition of an *α*-helical pore-forming toxin. The Journal of Physical Chemistry B 2016, 120, 12064–12078.

[10] Kulshrestha, A.; Punnathanam, S. N.; Ayappa, K. G. Finite temperature string method with umbrella sampling using path collective variables: application to secondary structure change in a protein. Soft Matter 2022, 18, 7593–7603.

[11] Matsuzaki, K.; Sugishita, K.; Fujii, N.; Miyajima, K. Molecular basis for membrane selectivity of an antimicrobial peptide, magainin 2. Biochemistry 1995, 34, 3423–3429.

[12] Tweten, R. K. Cholesterol-dependent cytolysins, a family of versatile pore-forming toxins. Infection and immunity 2005, 73, 6199–6209.

[13] Gilbert, R. J. Cholesterol-dependent cytolysins. Proteins Membrane Binding and Pore Formation 2010, 56–66.

[14] Oscarsson, J.; Mizunoe, Y.; Li, L.; Lai, X.-H.; Wieslander, Å.; Uhlin, B. E. Molecular analysis of the cytolytic protein ClyA (SheA) from Escherichia coli. Molecular Microbiology 1999, 32, 1226–1238.

[15] Sathyanarayana, P.; Visweswariah, S. S.; Ayappa, K. G. Mechanistic insights into pore Formation by an *α*-Pore forming toxin: Protein and lipid bilayer interactions of cytolysin A. Accounts of Chemical Research 2020, E7323–E7330.

[16] Sathyanarayana, P.; Maurya, S.; Behera, A.; Ravichandran, M.; Visweswariah, S. S.; Ayappa, K. G.; Roy, R. Cholesterol promotes cytolysin A activity by stabilizing the intermediates during pore formation. Proceedings of the National Academy of Sciences 2018, 115, E7323–E7330.

[17] Branduardi, D.; Gervasio, F. L.; Parrinello, M. From A to B in free energy space. The Journal of Chemical Physics 2007, 126, 054103.

[18] Tanaka, K.; Caaveiro, J. M.; Morante, K.; Gonźalez-Mañas, J. M.; Tsumoto, K. Structural basis for self-assembly of a cytolytic pore lined by protein and lipid. Nature Communications 2015, 6, 1–11.

[19] Varadarajan, V.; Desikan, R.; Ayappa, K. G. Assessing the extent of the structural and dynamic modulation of membrane lipids due to pore forming toxins: insights from molecular dynamics simulations. Soft Matter 2020, 16, 4840–4857.

[20] Desikan, R.; Maiti, P. K.; Ayappa, K. G. Assessing the structure and stability of transmembrane oligomeric intermediates of an *α*-helical toxin. Langmuir 2017, 33, 11496– 11510.

[21] Jo, S.; Kim, T.; Iyer, V. G.; Im, W. CHARMM-GUI: a web-based graphical user interface for CHARMM. Journal of Computational Chemistry 2008, 29, 1859–1865.

[22] Pronk, S.; Páll, S.; Schulz, R.; Larsson, P.; Bjelkmar, P.; Apostolov, R.; Shirts, M. R.; Smith, J. C.; Kasson, P. M.; van der Spoel, D., et al. GROMACS 4.5: a high-throughput and highly parallel open source molecular simulation toolkit. Bioinformatics 2013, 29, 845–854.

[23] Wang, J.; Wolf, R. M.; Caldwell, J. W.; Kollman, P. A.; Case, D. A. Development and testing of a general Amber force field. Journal of Computational Chemistry 2004, 25, 1157–1174.

[24] Jambeck, J. P.; Lyubartsev, A. P. Derivation and systematic validation of a refined all-atom force field for phosphatidylcholine lipids. The Journal of Physical Chemistry B 2012, 116, 3164–3179.

[25] Desikan, R.; Behera, A.; Maiti, P. K.; Ayappa, K. G. Methods in Enzymology; Elsevier, 2021; Vol. 649; pp 461–502.

[26] Hess, B.; Bekker, H.; Berendsen, H. J.; Fraaije, J. G. LINCS: a linear constraint solver for molecular simulations. Journal of Computational Chemistry 1997, 18, 1463–1472.

[27] Parrinello, M.; Rahman, A. Polymorphic transitions in single crystals: A new molecular dynamics method. Journal of Applied Physics 1981, 52, 7182–7190.

[28] Hedger, G.; Koldsø, H.; Chavent, M.; Siebold, C.; Rohatgi, R.; Sansom, M. S. Cholesterol interaction sites on the transmembrane domain of the hedgehog signal transducer and class FG protein-coupled receptor smoothened. Structure 2019, 27, 549–559.

[29] Koldsø, H.; Shorthouse, D.; Hélie, J.; Sansom, M. S. Lipid clustering correlates with membrane curvature as revealed by molecular simulations of complex lipid bilayers. PLoS computational biology 2014, 10, e1003911.

[30] Allouche, A.-R. Gabedit—A graphical user interface for computational chemistry softwares. Journal of Computational Chemistry 2011, 32, 174–182.

[31] Yesylevskyy, S.; Ramseyer, C. Determination of mean and Gaussian curvatures of highly curved asymmetric lipid bilayers: the case study of the influence of cholesterol on the membrane shape. Physical Chemistry Chemical Physics 2014, 16, 17052–17061.

[32] Agrawal, A.; Apoorva, K.; Ayappa, K. Transmembrane oligomeric intermediates of pore forming toxin Cytolysin A determine leakage kinetics. RSC Advances 2017, 7, 51750–51762.

[33] Vaidyanathan, M.; Sathyanarayana, P.; Maiti, P. K.; Visweswariah, S. S.; Ayappa, K. Lysis dynamics and membrane oligomerization pathways for Cytolysin A (ClyA) pore-forming toxin. RSC Advances 2014, 4, 4930–4942.

[34] Miller, C. M.; Brown, A. C.; Mittal, J. Disorder in cholesterol-binding functionality of CRAC peptides: A molecular dynamics study. The Journal of Physical Chemistry B 2014, 118, 13169–13174.

[35] Fantini, J.; Barrantes, F. J. How cholesterol interacts with membrane proteins: an exploration of cholesterol-binding sites including CRAC, CARC, and tilted domains. Frontiers in physiology 2013, 4, 31.

[36] Plesnar, E.; Subczynski, W. K.; Pasenkiewicz-Gierula, M. Saturation with cholesterol increases vertical order and smoothes the surface of the phosphatidylcholine bilayer: A molecular simulation study. Biochimica et Biophysica Acta (BBA)-Biomembranes 2012, 1818, 520–529.

[37] Ramakrishnan, N.; Bradley, R. P.; Tourdot, R. W.; Radhakrishnan, R. Biophysics of membrane curvature remodeling at molecular and mesoscopic lengthscales. Journal of Physics: Condensed Matter 2018, 30, 273001.

[38] Chakraborty, H.; Sengupta, D. Preface to special issue on protein-mediated membrane remodeling. The Journal of Membrane Biology 2022, 255, 633–635.

[39] Viswanathan, G.; Jafurulla, M.; Kumar, G. A.; Raghunand, T. R.; Chattopadhyay, A. Dissecting the membrane cholesterol requirement for mycobacterial entry into host cells. Chemistry and Physics of Lipids 2015, 189, 19–27.

