## Supporting Information for "Cholesterol catalyzes unfolding in membrane inserted motifs of the pore forming protein cytolysin A"

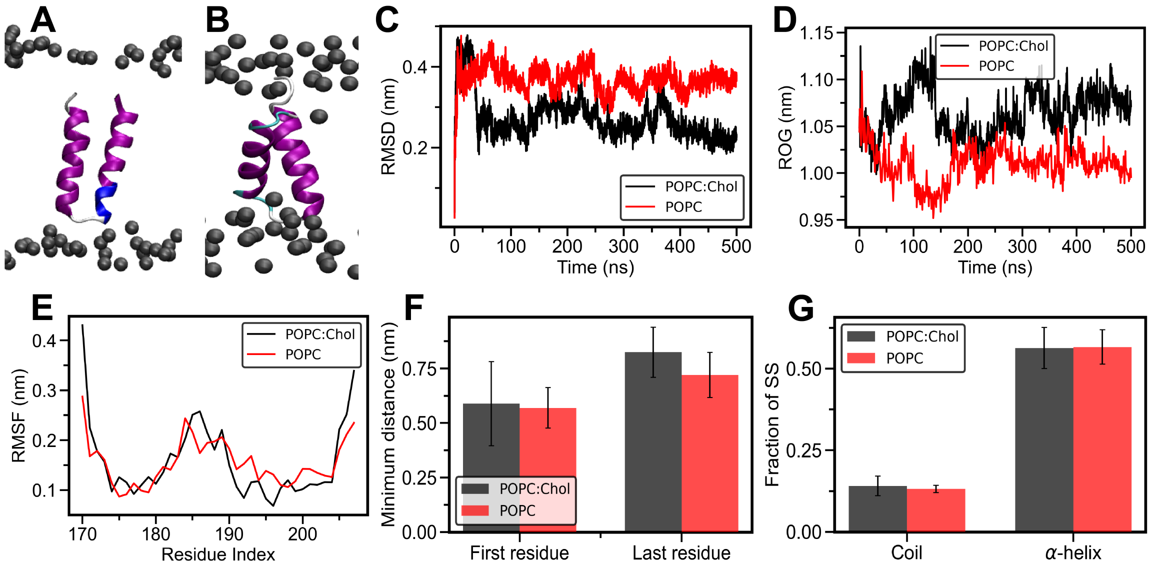


Figure 1: **Simulations of the helix-turn-helix motif of the protomer in membrane.** (A) Snapstho of initial protomer state in the POPC membrane. (B) The final snapshot of the protomer in the POPC membrane. Root mean squared deviation (RMSD), radius of gyratio (ROG) and root mean squared fluctuation (RMSF) comparison betwen POPC:Chol and POPC membrane in (C), (D), and (E), respectively. (F) The minimum distance between the C-$\alpha$ atom of the first and last residue to the lipid headgroups. (G) Fraction of secondary structure content.

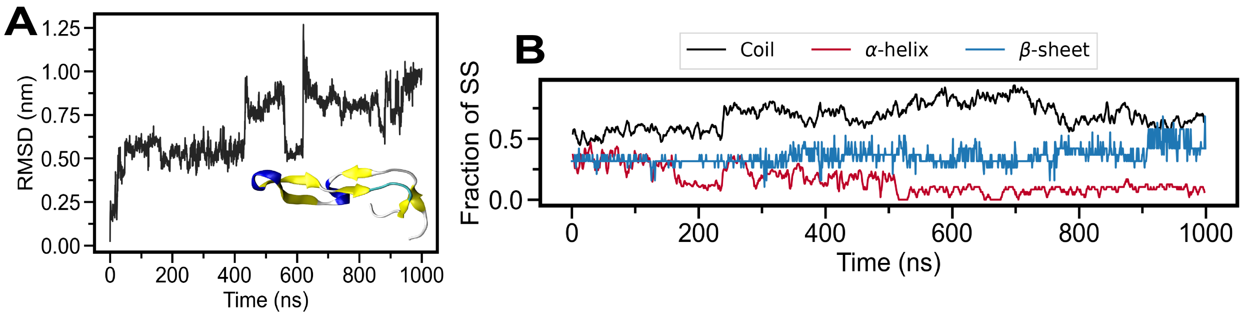


Figure 2: **Solvated $\beta$-tongue simulation.** (A) RMSD of the solvated $\beta$-tongue (170-207), inset is the final snapshot at 1 $\mu$s and (B) evolution of the secondary structure content with time.

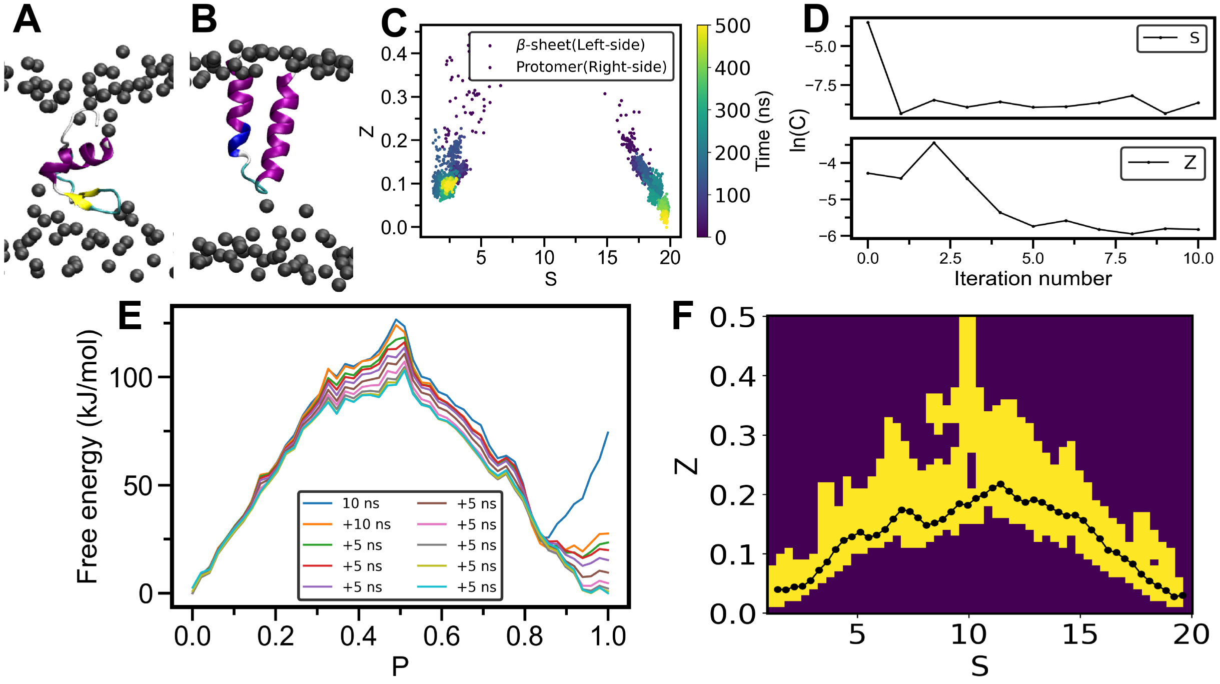


Figure 3: **String method detail For the membrane without cholesterol ($POPC$).** (A) Initial and (B) final end states, (C) unbiased sampling in $S$ and $Z$ space for the end states. (D) Convergence of the string, and (E) convergence of the free energy profile along the converged string. (E) Illustration of the number of times a specific grid point in the S-Z space is sampled during the umbrella sampling simulations over the entire simulation trajectory of ~3 microseconds. During the umbrella sampling, the $S$ and $Z$ space is divided into 50 equal bins resulting in a total of 2500 bins. Yellow color represents the bins with at least 0.1 million samples, while the darker regions represent regions where the numbers of points sampled are lower than 0.1 million. The regions around the vicinity of the converged path indicate adequate overlap of the histograms for the umbrella sampling simulations.

Table 1: **ClyA $\beta$-tongue simulation details.**

| Protein | Number of residues and (water) molecules | Ensemble NaCl concentration (M) | Number of POPC and (cholesterol) molecules | Simulation time (ns) |
|---|---|---|---|---|
| $\beta$-tongue exposed to membrane | 38 (8831) | NPT (1 atm, 310 K) 0.15 | 140 (60) | 1000 |
| $\beta$-tongue exposed to membrane | 38 (8831) | NPT (1 atm, 310 K) 0.15 | 140 (0) | 1000 |
| $\beta$-tongue placed in the membrane | 38 (8831) | NPT (1 atm, 310 K) 0.15 | 140 (60) | $500 \times 3$ |
| $\beta$-tongue placed in the membrane | 38 (8831) | NPT (1 atm, 310 K) 0.15 | 140 (0) | $500 \times 3$ |
| $\beta$-tongue in the solution | 38 (8831) | NPT (1 atm, 310 K) 0.15 | 0 (0) | 1000 |
| Protomer placed in membrane | 38 (8831) | NPT (1 atm, 310 K) 0.15 | 140 (60) | 500 |
| Protomer placed in membrane | 38 (8831) | NPT (1 atm, 310 K) 0.15 | 140 (0) | 500 |

Table 2: **Converged string location for POPC:Chol membrane.**

| Point | S | Z | Point | S | Z |
|---|---|---|---|---|---|
| 1 | 1.0396 | 0.0114 | 26 | 10.6419 | 0.1640 |
| 2 | 1.4238 | 0.0239 | 27 | 11.0262 | 0.1705 |
| 3 | 1.8080 | 0.0329 | 28 | 11.4105 | 0.16683 |
| 4 | 2.1922 | 0.0423 | 29 | 11.7948 | 0.1641 |
| 5 | 2.5764 | 0.0508 | 30 | 12.1791 | 0.1616 |
| 6 | 2.9604 | 0.0650 | 31 | 12.5634 | 0.1584 |
| 7 | 3.3447 | 0.0733 | 32 | 12.9477 | 0.1526 |
| 8 | 3.7289 | 0.0816 | 33 | 13.3319 | 0.1432 |
| 9 | 4.1132 | 0.0845 | 34 | 13.7162 | 0.1405 |
| 10 | 4.4975 | 0.0897 | 35 | 14.1005 | 0.1400 |
| 11 | 4.8818 | 0.0973 | 36 | 14.4847 | 0.1295 |
| 12 | 5.2661 | 0.0990 | 37 | 14.8687 | 0.1136 |
| 13 | 5.6504 | 0.1026 | 38 | 15.2528 | 0.1006 |
| 14 | 6.0346 | 0.1119 | 39 | 15.6369 | 0.1069 |
| 15 | 6.4188 | 0.1243 | 40 | 16.0212 | 0.1068 |
| 16 | 6.8022 | 0.1479 | 41 | 16.4055 | 0.1080 |
| 17 | 7.1848 | 0.1805 | 42 | 16.7895 | 0.0951 |
| 18 | 7.5690 | 0.1915 | 43 | 17.1737 | 0.0881 |
| 19 | 7.9528 | 0.1933 | 44 | 17.5580 | 0.0852 |
| 20 | 8.3361 | 0.1690 | 45 | 17.9423 | 0.0813 |
| 21 | 8.7204 | 0.1696 | 46 | 18.3267 | 0.0808 |
| 22 | 9.1047 | 0.1747 | 47 | 18.7110 | 0.0810 |
| 23 | 9.4890 | 0.1725 | 48 | 19.0953 | 0.0824 |
| 24 | 9.8733 | 0.1685 | 49 | 19.4790 | 0.0602 |
| 25 | 10.2576 | 0.1607 | 50 | 19.8623 | 0.0320 |

Table 3: **Converged string location for POPC membrane.**

| Point | S | Z | Point | S | Z |
|---|---|---|---|---|---|
| 1 | 1.42 | 0.04 | 26 | 10.6849 | 0.1995 |
| 2 | 1.7908 | 0.0391 | 27 | 11.0555 | 0.2108 |
| 3 | 2.1615 | 0.0441 | 28 | 11.4261 | 0.2173 |
| 4 | 2.5323 | 0.0451 | 29 | 11.7967 | 0.2054 |
| 5 | 2.903 | 0.0545 | 30 | 12.1672 | 0.1931 |
| 6 | 3.2734 | 0.0721 | 31 | 12.538 | 0.1866 |
| 7 | 3.6439 | 0.0859 | 32 | 12.9087 | 0.1907 |
| 8 | 4.014 | 0.1072 | 33 | 13.2794 | 0.1868 |
| 9 | 4.3844 | 0.1223 | 34 | 13.6501 | 0.1779 |
| 10 | 4.7552 | 0.1288 | 35 | 14.0209 | 0.17 |
| 11 | 5.1257 | 0.1362 | 36 | 14.3916 | 0.1641 |
| 12 | 5.4963 | 0.1273 | 37 | 14.7624 | 0.1652 |
| 13 | 5.8671 | 0.1315 | 38 | 15.1328 | 0.1562 |
| 14 | 6.2378 | 0.1409 | 39 | 15.5033 | 0.1404 |
| 15 | 6.6081 | 0.1583 | 40 | 15.8738 | 0.1254 |
| 16 | 6.9785 | 0.1741 | 41 | 16.244 | 0.1059 |
| 17 | 7.3493 | 0.1713 | 42 | 16.6148 | 0.1019 |
| 18 | 7.7198 | 0.1605 | 43 | 16.9855 | 0.0914 |
| 19 | 8.0902 | 0.1477 | 44 | 17.3562 | 0.083 |
| 20 | 8.461 | 0.1518 | 45 | 17.7266 | 0.0657 |
| 21 | 8.8317 | 0.1579 | 46 | 18.0972 | 0.0544 |
| 22 | 9.2023 | 0.1704 | 47 | 18.4679 | 0.0456 |
| 23 | 9.5728 | 0.1842 | 48 | 18.8386 | 0.0384 |
| 24 | 9.9436 | 0.1818 | 49 | 19.2092 | 0.0274 |
| 25 | 10.3142 | 0.1933 | 50 | 19.58 | 0.03 |

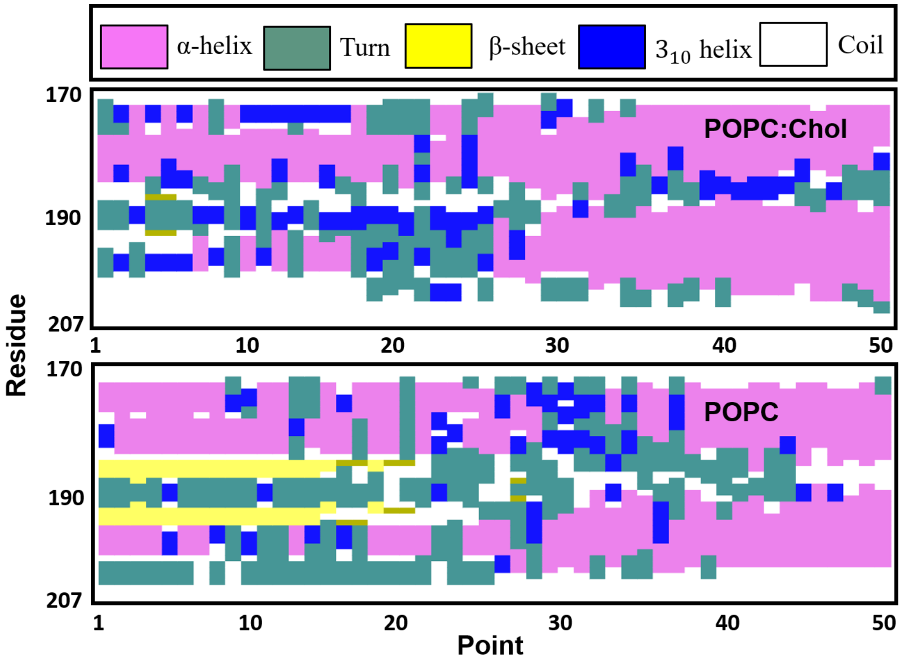


Figure 4: Comparing secondary structure changes on the path between POPC and POPC:Chol membrane. The structures on the path were taken from the end of the biased simulation for each point. The average values from the simulation is given in Figure 3C of the main manuscript.
